# Bayesian Inference of Bond Parameters from Single-Filament Interactions

**DOI:** 10.64898/2026.05.29.728470

**Authors:** Kristian A. T. Pajanonot, Sascha Lambert, Pallavi Kumari, Sarah Köster, Stefan Klumpp

## Abstract

We map interaction forces between two vimentin filaments (cytoskeletal components crucial for cell mechanics) using optical tweezers, while controlling the relative velocity. We introduce a powerful Bayesian inference framework to learn bond parameters directly from force trajectories. The information gained about the bond parameters is maximized by an optimal relative velocity and further by distributing measurements across multiple velocities. Our Bayesian framework is broadly applicable to a large range of biomolecular interactions and force spectroscopy techniques.

---

The mechanical response of biological cells to external forces is governed by the cytoskeleton, a composite network of microtubules, actin filaments, and intermediate filaments (IFs) such as vimentin.^1,2^ Its mechanical properties arise not only from the properties of individual filaments but also from the interactions between them, mediated through direct contacts^3–6^ and associated crosslinker proteins.^7–11^ Bonds between two individual filaments can be probed precisely with a quadruple optical trap setup.^5,6^ These experiments belong to a broader class of force spectroscopy experiment that exert force onto biomolecules to either probe interactions between two molecules or internal interactions within one molecule using different experimental techniques such as optical tweezers, magnetic tweezers, and atomic force microscopy (AFM).^12^

However, the quantitative analysis of the resulting force-time trajectories, aiming at inferring parameters of force-dependent bond rupture, is challenging despite the precision of the experimental methods, as bond rupture is inherently stochastic and loading rate dependent. The force-time trajectory before bond rupture typically depends on the details of the experimental setup, as it reflects the stretching of the molecules of interest themselves or of polymeric linkers. Therefore, their analysis requires to model how the force on the bond is built up. Existing bond parameter estimation methods typically rely on reduced representations of force spectroscopy data. In AFM experiments with multiple loading rates, parameters are obtained from a linear fit of the most probable rupture force as a function of the logarithm of the loading rate. ^13–15^ Extensions of this framework fit the full rupture force distribution rather than only its peak,^16,17^ extracting additional information about the underlying energy landscape. When only a single loading rate is available, simulated and experimental rupture force distributions are compared via a Kolmogorov–Smirnov test.^5,6,18^ However, these approaches discard the information contained in the force trajectory before rupture and do not provide full uncertainty quantification.

Here, we propose a Bayesian inference framework that uses the full force trajectory, yielding a posterior distribution over bond parameters. Related Bayesian approaches are also increasingly adopted in the analysis of single molecule fluorescence experiments,^19–21^ which shares some of the challenges, although not those related to the time-dependent force. An advantage of our Bayesian framework is that it is applicable regardless of whether experiments are performed at single or multiple loading rates, and that it does not require a model of the time evolution of the force on the bond, but uses the experimental data directly. Moreover, it directly quantifies parameter uncertainty. We validate our framework against existing vimentin–vimentin interaction data,^5^ demonstrating improved parameter estimation over existing approaches. We further show that by performing quadruple optical trap experiments across multiple pulling velocities, the framework can be used to guide experimental design, identifying the conditions that most efficiently constrain the bond parameters.

## Interactions between two vimentin intermediate filaments

We study the interaction between two single vimentin IFs using a quadruple optical trap experiment^5,6^ (see Fig. S1 in the Supplemental Material^22^ for a schematic of the experimental setup and Sec. I for a detailed description of the experimental methods). In this experiment, two filaments are held with four optical traps positioned in a cross configuration and brought into contact to allow bond formation (Fig. 1a). When the “vertical” filament (oriented along the *y*-direction) is moved in the *y*-direction at a fixed pulling velocity *v*, the two filaments eventually interact and form a bond, bending the “horizontal” filament (oriented along the *x* direction, Fig. 1b(ii)). Continued pulling on the vertical vimentin IF increases the load on the bond until rupture occurs, after which the horizontal filament returns to its initial straight configuration (Fig. 1b(iii)). To measure the force acting on the horizontal vimentin IF, we record the force *F*_1*y*_ on trap 1 (Fig. 1b) and calculate the total force *F* on the bond based on the geometric arrangement of the filaments derived from the bead positions (see Supplemental Material^22^ Fig. S2 and Sec. II for a detailed description of the data analysis). Fig. 1c shows representative force-time data of an interaction event where we can assign the measured force to two distinct bond states: a bound state and an unbound state. In the bound state the two filaments interact and the force increases (red scatter). The unbound state occurs immediately after when the interaction breaks at the breaking force *F*_*B*_ and the force abruptly drops to 0 pN (blue scatter). This classification reflects how we model the interaction as a single bond with two states, (Fig. 1d), *i*.*e*., the simplest model consistent with the observation that each interaction event shows a single continuous and monotonous force increase followed by an abrupt rupture. We use Bell-Evans kinetics^13,23,24^ with the force-dependent unbinding rate

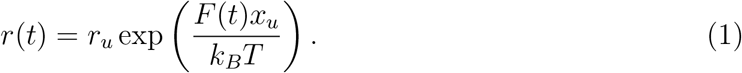

**Figure 1.**
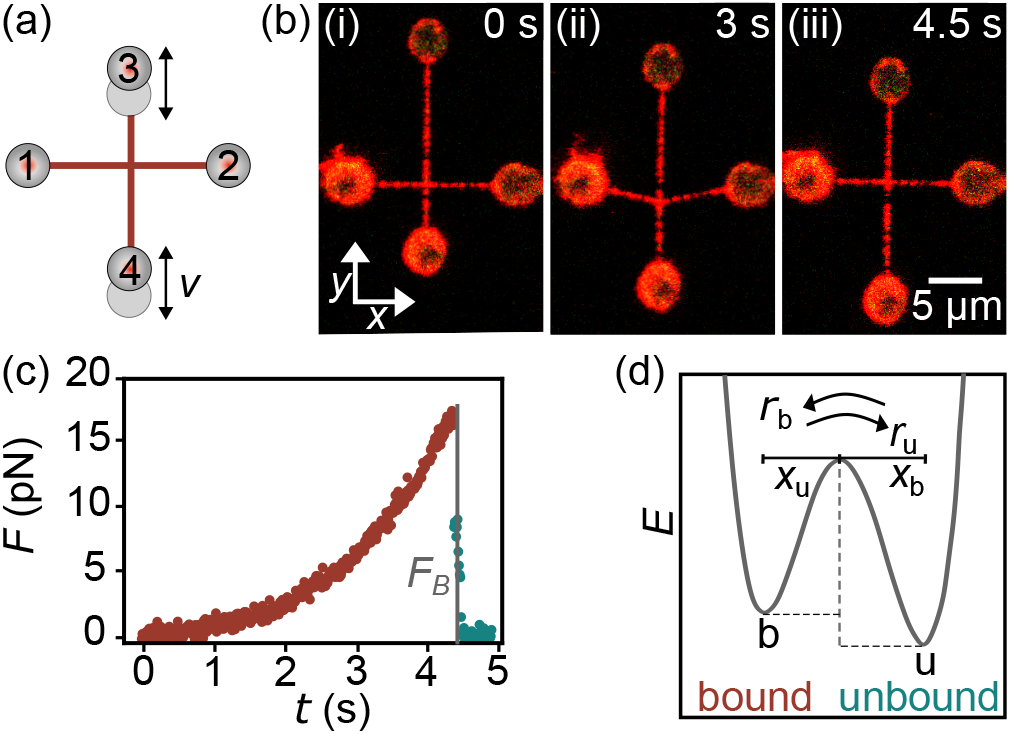
(a) Sketch of two vimentin IFs held between optically trapped beads in a crossed configuration allowing for bond formation and measurement of their interaction forces by pulling the vertical filament at a velocity *v*. (b) Confocal image series showing an interaction event where the horizontal filament bends and then returns to its initial conformation after the interaction breaks. (c) Typical force-time data of an interaction event. Once the filaments interact, the force increases until the bond breaks at the breaking force *F*_*B*_, where *F* abruptly drops to 0 pN. The red and blue scatter denote the bound and unbound states, respectively. (d) Energy landscape for theoretically modeling of the interaction as a single bond characterized by the force-independent unbinding rate *r*_*u*_ and the force sensitivity of the bond *x*_*u*_.

Here *r*_*u*_ is the force-free unbinding rate, and *x*_*u*_ is the distance from the bound state to the transition state interpreted as the force sensitivity of the bond. These two parameters jointly govern the observed rupture behavior which cannot be measured directly from experiments and must be inferred from force-time trajectories.

## Bayesian inference

To determine the bond parameters *r*_*u*_ and *x*_*u*_, we develop a Bayesian inference framework for quantifying how plausible different parameter combinations are given the measured force trajectories. Each sampled time point *i* in a force trajectory provides a pair of observables: the measured force *F*_*i*_ and the corresponding bond state *S*_*i*_ ∈ {0, 1}, where *S*_*i*_ = 1 denotes a bound event and *S*_*i*_ = 0 an unbound/rupture event. All such {*F*_*i*_, *S*_*i*_} pairs from all binding events (from the first bound state observation to the first unbound one) across all trajectories in a given experimental condition are pooled into a dataset for that condition that is used to constrain the two bond parameters *r*_*u*_ and *x*_*u*_. Frequent bond rupture at relatively low forces suggests a higher intrinsic unbinding rate *r*_*u*_, whereas differences in bond survival between low and high forces reflect the degree of force sensitivity of the bond, characterized by *x*_*u*_. We aim to compute the posterior *P* (*r*_*u*_, *x*_*u*_|{*F*_*i*_, *S*_*i*_}), which captures both the parameter uncertainties and the correlations between the parameters. The posterior is obtained via Bayes’ rule:

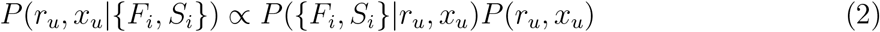

with uniform priors *P* (*r*_*u*_, *x*_*u*_) over plausible ranges much wider than the resulting posteriors. The likelihood *P* ({*F*_*i*_, *S*_*i*_}|*r*_*u*_, *x*_*u*_) is constructed from the pair of *F*_*i*_, *S*_*i*_ observations at discrete time intervals Δ*t* = *t*_*i*+1_−*t*_*i*_, over which the force is assumed constant. We assume Markovian dynamics, under which the bond has no memory and survival follows an exponential decay process, so the probability of remaining bound over Δ*t* depends only on the instantaneous detachment rate, not on how long the bond has been intact. Each observation therefore contributes a Bernoulli likelihood term:

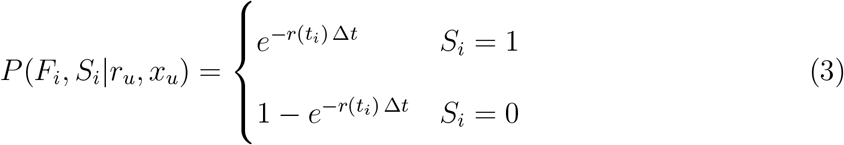

where the two cases correspond to bond survival and unbinding, respectively. Since observations at each time step are independent under the Markov assumption, the full likelihood is the product over all data points:

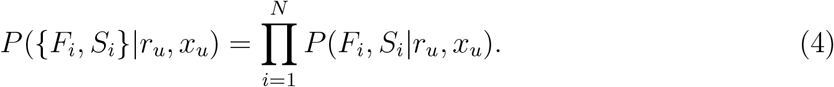

Having the prior and likelihood, we estimate the posterior numerically using Markov Chain Monte Carlo (MCMC) sampling via the PyMC package, ^25,26^ employing the No-U-Turn Sampler (NUTS)^27^ which efficiently explores the *r*_*u*_, *x*_*u*_ parameter space (see Supplemental Material^22^ Sec. IIIA-D for details on the implementation of the Bayesian inference framework).

## Validation of the Bayesian framework

To validate the Bayesian inference framework, we use it to reanalyze published experimental data of vimentin-vimentin interactions. ^5^ In this study, Schepers et al. investigated the interaction between two single vimentin IFs in different buffer conditions to determine whether the interactions are driven by electrostatic or hydrophobic forces, or both, and recorded the experimental distributions of the breaking force *F*_*B*_ for each buffer condition. To identify valid parameter pairs (*r*_*u*_, *x*_*u*_), Monte Carlo simulations of rupture events were performed for each candidate pair, and the resulting simulated *F*_*B*_ distribution was compared to the experimental one using the Kolmogorov-Smirnov (KS) test,^28^ which quantifies whether two distributions could plausibly have been drawn from the same underlying distribution. Valid parameter pairs were identified as the 95% confidence region, shown as the uniformly colored regions in Fig. 2.

**Figure 2.**
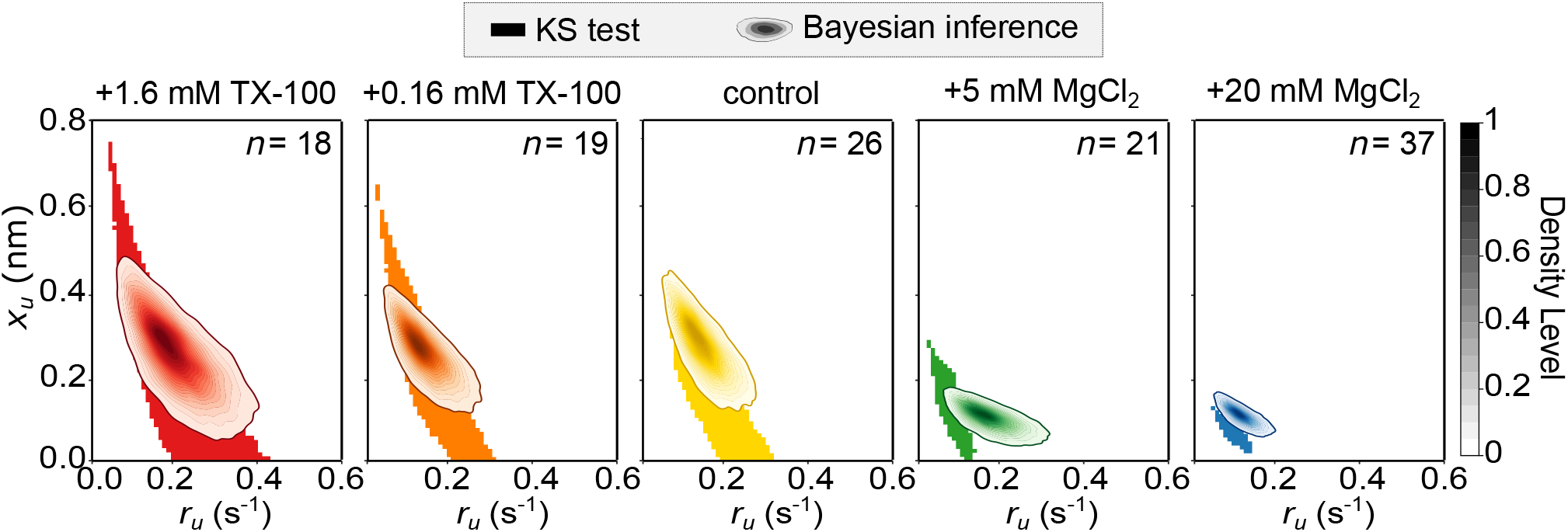
Comparison of bond parameter estimates obtained using a Kolmogorov–Smirnov (KS) test and Bayesian inference on previously published vimentin-vimentin interaction data ^5^ acquired in different buffer conditions. The Bayesian framework shows probability distributions (95% credible interval), where darker regions indicate bond parameters with higher probability. The Bayesian inference results overlap with the regions accepted by the KS test and indeed yield narrower parameter estimates.

However, this approach has several disadvantages: It reduces each force trajectory to a single summary statistic, *F*_*B*_, and discards all information about the force evolution prior to rupture; valid parameter pairs are identified only indirectly through Monte Carlo simulations of rupture events, which require additional model assumptions; and it provides only a binary acceptance region. In contrast, our Bayesian framework directly incorporates force and bond state information at every sampled time point along each trajectory and, thus, utilizes the full experimental data, requires no simulation and provides a posterior probability distribution over (*r*_*u*_, *x*_*u*_) offering quantification of parameter uncertainty.

Figure 2 compares the two approaches across all buffer conditions. The Bayesian posteriors (shown as 95% credible intervals with darker regions indicating parameter combinations with higher probability density) overlap with the KS-accepted regions in all conditions, and the trends reported in Ref.^5^ such as the decrease in *r*_*u*_ and *x*_*u*_ under MgCl_2_ conditions are consistently recovered. We observe minor discrepancies, the decrease in *r*_*u*_ is consistent between the two methods for 20 mM MgCl_2_ but less pronounced in the 5 mM MgCl_2_ condition in the Bayesian results. This can be understood from how each method uses the data. Despite these minor discrepancies, there is an overall good agreement between the two methods which therefore supports the validity of the Bayesian framework.

## Optimal experimental design by combining multiple pulling velocities

stHaving validated the framework, we next investigate how the pulling velocity *v* can be exploited to vary the information we gain about the bond parameters. The magnitude of the breaking force *F*_*B*_ is set by how fast the vertical filament is pulled. Slower pulling mostly results in unbinding at low *F*_*B*_, with a rate near the intrinsic unbinding rate *r*_*u*_, dominated by thermal fluctuations, while faster pulling builds force rapidly, making the force-dependent exponential term in Eq. 1 increasingly dominant. Varying the pulling velocity therefore probes different regimes of bond kinetics. We thus perform vimentin-vimentin experiments across a range of pulling velocities *v* from 0.05 to 5 *µ*m/s (representative force-time curves are shown in Fig. S3 in Supplemental Material^22^). Figure 3a shows that increasing *v* shifts the breaking force distribution toward higher *F*_*B*_ and broadens it. Before analyzing the posteriors, we note that at higher *v*, we observe events with rebinding under force (see Fig. S4a in Supplemental Material^22^) where the bond ruptures and the force drops to non-zero force, but then the system rebinds, causing the force to rise again. These arise from new contact points forming along the filament as it is pulled, rather than reformation of the original bond, and occur more frequently at higher *v* (Supplemental Material, ^22^ Fig. S4b). Since these rebinding events are too few to allow reliable parameter estimation, and the resulting posteriors are different from those obtained from single-binding events (Supplemental Material,^22^ Fig. S4c), they are excluded from the following analysis.

**Figure 3.**
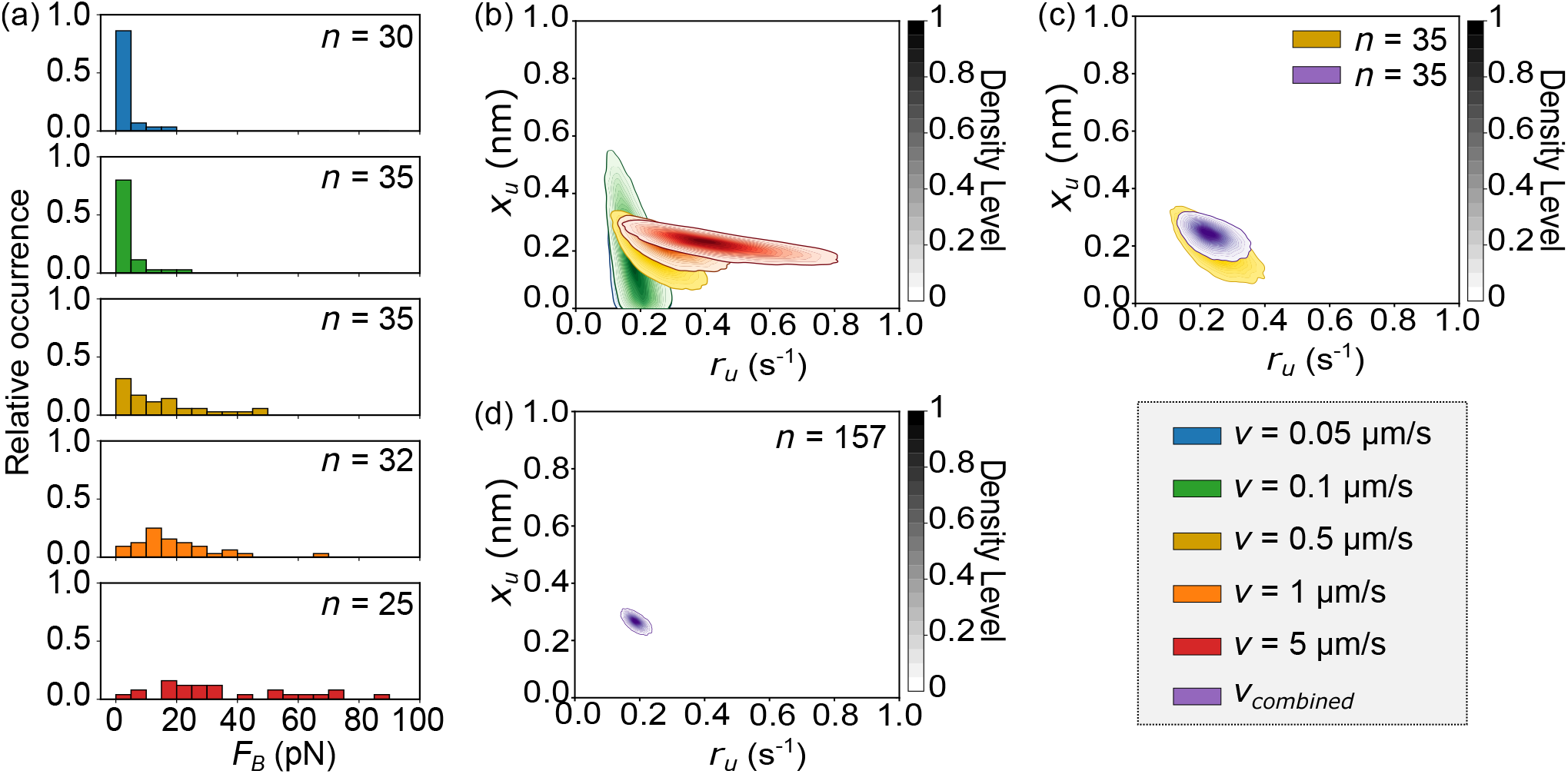
Improved bond parameter estimation using different pulling velocities, *v*. (a) Breaking force (*F*_*B*_) distributions widen with increasing *v*. (b) Bayesian inference results reveal an overlap of posteriors across different *v*. Low *v* better constrain *r*_*u*_, whereas high *v* better constrain *x*_*u*_. (c) Posterior distributions from the same number of events (*n* = 35), comparing single-velocity (shown in yellow) versus combined velocities (*v* between 0.05 and 5 *µ*m/s, shown in purple). This comparison clearly shows that the posterior narrows for combined *v*. (d) Increasing *n* further narrows the posterior, illustrating improved precision with more data.

When we apply Bayesian inference for each pulling velocity *v* as shown in Fig. 3b, the resulting posteriors overlap across all pulling velocities. We expect this overlap, as *r*_*u*_ and *x*_*u*_ are intrinsic properties of the bond that are independent of how the force is applied, and should therefore be consistently recovered across different pulling velocities. The posteriors in Fig. 3b also reveal that different pulling *v* constrain different bond parameters. Low *v* gives a narrow posterior for *r*_*u*_ (*r*_*u*_ = 0.18 ± 0.04 s^−1^) but a broad posterior for *x*_*u*_ (*x*_*u*_ = 0.13 ± 0.11 nm) (see Fig. S5a, Supplemental Material^22^), because at low force Eq. 1 is dominated by the force-independent term *r*_*u*_. In contrast, high *v* gives a posterior with narrow *x*_*u*_ (*x*_*u*_ = 0.22 ± 0.03 nm) but a broad posterior for *r*_*u*_ (*r*_*u*_ = 0.45 ± 0.14 s^−1^) (see Fig. S5e, Supplemental Material^22^), because at high force the exponential term with *x*_*u*_ dominates Eq. 1. This trade-off suggests that combining data across velocities should yield sharper parameter estimates than any single *v* alone. Indeed, Fig. 3c shows that combining (*n* = 7) events from each *v* yields a sharper posterior than a single *v* dataset of the same total size (*n* = 35), demonstrating that distributing measurements across multiple pulling velocities is a more efficient experimental design than collecting many repeats at a single velocity. Combining all available data sets (*n* = 157) spanning the full range of *v* yields a highly precise joint posterior (Fig. 3d), with both parameters simultaneously well-constrained due to the velocity-dependent sensitivity to *r*_*u*_ and *x*_*u*_.

## Maximizing the information gain

A quantitative measure of the information that we gain about the bond parameters upon updating from prior to posterior is provided by the Kullback-Leibler (KL) divergence:^29^

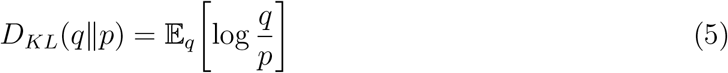

where *q* = *P* (*r*_*u*_, *x*_*u*_|{*F*_*i*_, *S*_*i*_}) denotes the posterior and *p* = *P* (*r*_*u*_, *x*_*u*_) the prior. A higher value of *D*_*KL*_ indicates that there is a large difference between the posterior and prior, meaning the data have strongly narrowed down the plausible parameter values. Figure 4a shows the information gain for *r*_*u*_ and *x*_*u*_ at each pulling velocity *v* for *n* = 25 events per condition, confirming the velocity-dependent sensitivity seen in the posteriors. The information gain for *r*_*u*_ (blue) decreases with *v* while the information gain for *x*_*u*_ (red) increases with *v*. Figure 4b shows the information gain in the joint posterior over (*r*_*u*_, *x*_*u*_). Among individual velocities, *v* = 1 *µ*m/s yields the highest information gain of ∼4.7 bits identifying this as the optimal pulling velocity in the single-velocity case. Combining data across multiple velocities (dashed lines, *n* = 25) further increases the joint information gain to ∼5.7 bits, demonstrating that integrating measurements over multiple pulling velocities *v* outperforms any single-velocity approach (see Supplemental Material^22^ Sec. IIIE for details on the quantification of *D*_*KL*_).

**Figure 4.**
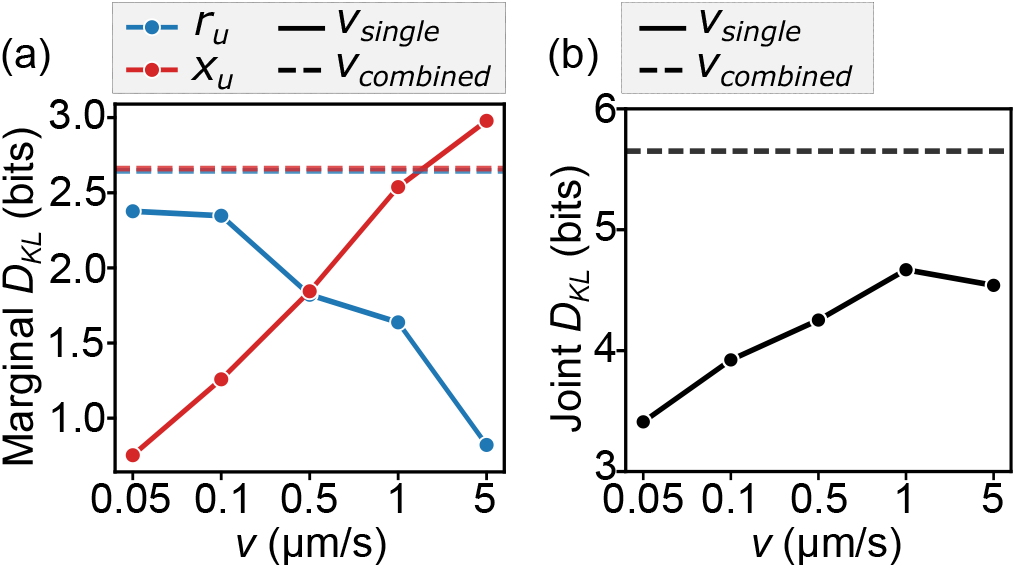
(a) Marginal information gain, quantified as the KL divergence (*D*_*KL*_) from prior to posterior, for *r*_*u*_ (blue) and *x*_*u*_ (red) individually across pulling velocities *v* (solid lines, *n* = 25 events per condition). The information gain for *r*_*u*_ is highest at *v* = 0.05 *µ*m/s and decreases with *v*, while *x*_*u*_ shows the opposite trend, reflecting the velocity-dependent sensitivity trade-off between the two parameters. Combining data across all velocities yields high information gain for both parameters simultaneously (dashed lines). (b) Joint information gain, quantified as the *D*_*KL*_ of the joint posterior over (*r*_*u*_, *x*_*u*_), across pulling velocities *v* (*n* = 25 events per condition). Among single velocities, *v* = 1 *µ*m/s yields the highest joint information gain (∼4.7), while combining data across all velocities increases this to ∼5.7, outperforming any single-velocity strategy.

## Concluding remarks

We present a Bayesian inference framework to estimate bond parameters from force-time curves in force spectroscopy experiments. While we apply it here to the interaction of two vimentin IFs, the method is general and can also be applied to data obtained with other experimental techniques such as AFM^13–15^ and to bonds between other (biomolecular) binding partners. As an example we apply the approach to the interaction between a vimentin IF and a microtuble,^6^ shown in Fig. S6 in Supplemental Material. ^22^

The Bayesian approach provides a posterior probability distribution of the parameters that also represents their uncertainty and correlations between them. Importantly, the method makes use of the full force-time curve rather than just the breaking force and does not require a model for how the force increases with time but rather uses the experimental time series of the force directly. Only a model for the force-dependent rupture of the bond of interest is required. In our case, this is given by Bell-Evans kinetics, but the method can equally be applied to other models of force-dependent unbinding. Extending our current approach, competing mechanistic descriptions can be distinguished with Bayesian model comparison.^19^ Beyond parameter estimation, the Bayesian framework serves as a guide for optimal experimental design, maximizing the information gained from a limited number of experiments, in our case by combining measurements across different velocities. Moreover, if the posterior reveals that one parameter remains poorly constrained, the framework itself indicates which pulling velocity would be most informative to pursue next, providing a basis for iterative data collection.

In conclusion, the Bayesian inference framework introduced here provides a tool for the quantitative analysis of force spectroscopy experiment and enables data-driven experimental design. We expect that it can be applied and generalized broadly, including to different experimental methods, different force-dependencies (e.g., slip and catch bonds) as well as different interactions including those between two biomolecules, intramolecular interactions and interactions mediated by multiple bonds.

## Acknowledgement

We thank Pratima Sawant, Komal Bhattacharyya, Anna Schepers, and Charlotta Lorenz for helpful discussions, and Kamila Sabagh and Anna Dudek for technical support. The authors acknowledge funding by the Deutsche Forschungsgemeinschaft (DFG, German Research Foundation) – Project-ID 449750155 – RTG 2756, Projects A3 (to S.Kl.) and A2, A7 (to S.Kö.). This research was conducted within the Max Planck School Matter to Life, supported by the Dieter Schwarz Foundation and the German Federal Ministry of Research, Technology and Space (BMFTR) in collaboration with the Max Planck Society (K.A.T.P., S.Kö., S.Kl.). P.K. acknowledges funding by the European Union’s Horizon Europe program under the Marie Skłodowska-Curie Actions (MSCA), Grant No. 101148781.

## References

(1) Fletcher, D. A.; Mullins, R. D. Cell mechanics and the cytoskeleton. Nature 2010, 463, 485–492.

(2) Lorenz, C.; Köster, S. Multiscale architecture: Mechanics of composite cytoskeletal networks. Biophysics Reviews 2022, 3, 031304.

(3) Huber, F.; Boire, A.; López, M. P.; Koenderink, G. H. Cytoskeletal crosstalk: When three different personalities team up. Current Opinion in Cell Biology 2015, 32, 39–47.

(4) Inoue, D.; Obino, D.; Pineau, J.; Farina, F.; Gaillard, J.; Guerin, C.; Blanchoin, L.; Lennon-Duménil, A.; Théry, M. Actin filaments regulate microtubule growth at the centrosome. The EMBO Journal 2019, 38, e99630.

(5) Schepers, A. V.; Lorenz, C.; Nietmann, P.; Janshoff, A.; Klumpp, S.; Köster, S. Multiscale mechanics and temporal evolution of vimentin intermediate filament networks. Proceedings of the National Academy of Sciences 2021, 118, e2102026118.

(6) Schaedel, L.; Lorenz, C.; Schepers, A. V.; Klumpp, S.; Köster, S. Vimentin intermediate filaments stabilize dynamic microtubules by direct interactions. Nature Communications 2021, 12, 3799.

(7) Mohan, R.; John, A. Microtubule-associated proteins as direct crosslinkers of actin filaments and microtubules. IUBMB Life 2015, 67, 395–403.

(8) Marks, P. C.; Hewitt, B. R.; Baird, M. A.; Wiche, G.; Petrie, R. J. Plectin linkages are mechanosensitive and required for the nuclear piston mechanism of three-dimensional cell migration. Molecular Biology of the Cell 2022, 33, ar104.

(9) Prechova, M.; Adamova, Z.; Schweizer, A.-L.; Maninova, M.; Bauer, A.; Kah, D.; Meier-Menches, S. M.; Wiche, G.; Fabry, B.; Gregor, M. Plectin-mediated cytoskeletal crosstalk controls cell tension and cohesion in epithelial sheets. Journal of Cell Biology 2022, 221, e202105146.

(10) Petitjean, I. I.; Tran, Q. D.; Goutou, A.; Kabir, Z.; Wiche, G.; Leduc, C.; Koenderink, G. H. Reconstitution of cytolinker-mediated crosstalk between actin and vimentin. European Journal of Cell Biology 2024, 103, 151403.

(11) Bareja, I.; Kučera, O.; Petitjean, I. I.; Orozco Monroy, B. E.; Sabo, J.; Braun, M.; Lansky, Z.; Koenderink, G. H.; Dogterom, M. Anillin directly crosslinks microtubules with actin filaments. The EMBO Journal 2025, 44, 4803–4824.

(12) Neuman, K. C.; Nagy, A. Single-molecule force spectroscopy: optical tweezers, magnetic tweezers and atomic force microscopy. Nature Methods 2008, 5, 491–505.

(13) Merkel, R.; Nassoy, P.; Leung, A.; Ritchie, K.; Evans, E. Energy landscapes of receptor–ligand bonds explored with dynamic force spectroscopy. Nature 1999, 397, 50–53.

(14) Verdorfer, T.; Gaub, H. E. Ligand Binding Stabilizes Cellulosomal Cohesins as Revealed by AFM-based Single-Molecule Force Spectroscopy. Scientific Reports 2018, 8, 9634.

(15) Goktas, M.; Luo, C.; Sullan, R. M. A.; Bergues-Pupo, A. E.; Lipowsky, R.; Vila Verde, A.; Blank, K. G. Molecular mechanics of coiled coils loaded in the shear geometry. Chemical Science 2018, 9, 4610–4621.

(16) Dudko, O.; Hummer, G.; Szabo, A. Intrinsic Rates and Activation Free Energies from Single-Molecule Pulling Experiments. Physical Review Letters 2006, 96, 108101.

(17) Dudko, O. K.; Hummer, G.; Szabo, A. Theory, analysis, and interpretation of single-molecule force spectroscopy experiments. Proceedings of the National Academy of Sciences 2008, 105, 15755–15760.

(18) Gennerich, A., Ed. Optical Tweezers: Methods and Protocols; Methods in Molecular Biology; Springer US: New York, NY, 2022; Vol. 2478; Chapter 25.

(19) Kinz-Thompson, C. D.; Ray, K. K.; Gonzalez, R. L. Bayesian Inference: The Comprehensive Approach to Analyzing Single-Molecule Experiments. Annual Review of Biophysics 2021, 50, 191–208.

(20) Bryan, J. S.; Tashev, S. A.; Fazel, M.; Scheckenbach, M.; Tinnefeld, P.; Herten, D.-P.; Pressé, S. Bayesian Inference of Binding Kinetics from Fluorescence Time Series. The Journal of Physical Chemistry B 2025, 129, 4670–4681.

(21) Mattamira, C.; Ward, A.; Krishnan, S. T.; Lamichhane, R.; Barrera, F. N.; Sgouralis, I. Bayesian analysis and efficient algorithms for single-molecule fluorescence data and step counting. Biophysical Journal 2025, 124, 3161–3173.

(22) See Supplemental Material [url], which includes Refs.,5,6,23–27,29–40 for details on the experimental methods, data analysis and Bayesian inference.

(23) Bell, G. I. Models for the Specific Adhesion of Cells to Cells: A theoretical framework for adhesion mediated by reversible bonds between cell surface molecules. Science 1978, 200, 618–627.

(24) Evans, E.; Ritchie, K. Dynamic strength of molecular adhesion bonds. Biophysical Journal 1997, 72, 1541–1555.

(25) Salvatier, J.; Wiecki, T. V.; Fonnesbeck, C. Probabilistic programming in Python using PyMC3. PeerJ Computer Science 2016, 2, e55.

(26) Abril-Pla, O.; Andreani, V.; Carroll, C.; Dong, L.; Fonnesbeck, C. J.; Kochurov, M.; Kumar, R.; Lao, J.; Luhmann, C. C.; Martin, O. A.; Osthege, M.; Vieira, R.; Wiecki, T.; Zinkov, R. PyMC: a modern, and comprehensive probabilistic programming framework in Python. PeerJ. Computer Science 2023, 9, e1516.

(27) Hoffman, M. D.; Gelman, A. The No-U-Turn Sampler: Adaptively Setting Path Lengths in Hamiltonian Monte Carlo. Journal of Machine Learning Research 2014, 15, 1593–1623.

(28) Massey, F. J. The Kolmogorov-Smirnov Test for Goodness of Fit. Journal of the American Statistical Association 1951, 46, 68–78.

(29) Cover, T. M.; Thomas, J. A. Elements of information theory; Wiley series in telecommunications; Wiley: New York, 1991; Chapter 2.

(30) Herrmann, H.; Kreplak, L.; Aebi, U. Methods in Cell Biology ; Elsevier, 2004; Vol. 78; pp 3–24.

(31) Forsting, J.; Kraxner, J.; Witt, H.; Janshoff, A.; Köster, S. Vimentin Intermediate Filaments Undergo Irreversible Conformational Changes during Cyclic Loading. Nano Letters 2019, 19, 7349–7356.

(32) Duane, S.; Kennedy, A.; Pendleton, B. J.; Roweth, D. Hybrid Monte Carlo. Physics Letters B 1987, 195, 216–222.

(33) Gelman, A.; Rubin, D. B. Inference from Iterative Simulation Using Multiple Sequences. Statistical Science 1992, 7, 457–472.

(34) Gelman, A.; Carlin, J. B.; Stern, H. S.; Dunson, D. B.; Vehtari, A.; Rubin, D. B. Bayesian Data Analysis; Chapman and Hall/CRC, 2013.

(35) Vanlier, J.; Pauszek, R.; Moldovan, D.; van den Berg, A.; Jachowski, T.; Mirone, A.; Broekmans, O.; Moerland, R.; Moyo, A.; lilfer Lamerton, S. lumicks/pylake: v1.8.0. 2025; 10.5281/zenodo.4280788.

(36) Janissen, R.; Berghuis, B. A.; Dulin, D.; Wink, M.; Van Laar, T.; Dekker, N. H. Invincible DNA tethers: Covalent DNA anchoring for enhanced temporal and force stability in magnetic tweezers experiments. Nucleic Acids Research 2014, 42, e137.

(37) Block, J.; Witt, H.; Candelli, A.; Danes, J. C.; Peterman, E. J. G.; Wuite, G. J. L.; Janshoff, A.; Köster, S. Viscoelastic properties of vimentin originate from nonequilibrium conformational changes. Science Advances 2018, 4, eaat1161.

(38) Winheim, S.; Hieb, A. R.; Silbermann, M.; Surmann, E. M.; Wedig, T.; Herrmann, H.; Langowski, J.; Mücke, N. Deconstructing the late phase of vimentin assembly by total internal reflection fluorescence microscopy (TIRFM). PLoS ONE 2011, 6, e19202–e19202.

(39) Nöding, B.; Köster, S. Intermediate filaments in small configuration spaces. Physical Review Letters 2012, 108, 088101.

(40) Block, J.; Witt, H.; Candelli, A.; Peterman, E. J.; Wuite, G. J.; Janshoff, A.; Köster, S. Nonlinear Loading-Rate-Dependent Force Response of Individual Vimentin Intermediate Filaments to Applied Strain. Physical Review Letters 2017, 118, 048101.

