## Supporting Material for "Bayesian Inference of Bond Parameters from Single-Filament Interactions"

### Supplemental Material for: Bayesian Inference of Bond Parameters from Single-Filament Interactions

### Materials and Methods

#### Protein preparation

The preparation of vimentin follows the protocol described in Refs.[1,2] Briefly, human vimentin C328N with additional amino acids GGC at the C terminus is purified from inclusion bodies. The purified protein is kept under denaturing conditions and is stored at -80°C. Fluorescent labeling with Atto647N-maleimide (Atto-Tech GmbH, Siegen, Germany) is described in the next section. Labeled and unlabeled protein are mixed to achieve a final degree of labeling 4%. Reconstitution is performed at room temperature by stepwise dialysis using a 50 kDa MWCO dialysis tubing (Dialysis membrane Spectra/Por 7, Carl Roth, Karlsruhe, Germany) against 2 mM phosphate buffer (PB) pH 7.5. The urea concentration is gradually reduced in 30-min intervals through sequential dialysis at 6 M, 4 M, 2 M, 1 M, and 0 M, followed by an overnight dialysis at 4°C. Protein concentration is measured by UV absorption spectroscopy (Nanodrop ND-1000, ThermoScientific Technologies, Inc., Wilmington, DE, USA) using an extinction coefficient of $\varepsilon_{\text{protein}}(280\,\text{nm}) = 24.87 \times 10^3$ M$^{-1}$ cm$^{-1}$ and a molecular mass of 53.88 kDa. For filament assembly, the reconstituted protein is dialyzed against assembly buffer (2 mM PB, 100 mM KCl, pH 7.5) at 36°C for 18 h at a protein concentration of 0.2 g/L. For optical tweezer measurements, vimentin is diluted in assembly buffer to obtain a final concentration of 1 mg/L.

#### Fluorescent labeling of vimentin

Vimentin is labeled following the protocol described in Refs.[3–5] The protein previously stored in -80°C is thawed and dialyzed overnight using 50 kDa MWCO dialysis tubing against the labeling buffer (50 mM PB with 5 M urea at pH 7.0) at 4°C. After dialysis, the protein concentration is adjusted to 1 g/L using UV absorption spectroscopy. Atto647N-maleimide is first dissolved in DMSO at a concentration of 10 mM. The dye is then added to 1 mL of protein in a stepwise manner with four additions of 5 $\mu$L each to reach a final concentration of 0.2 mM. After each addition, the mixture is placed on a shaker at 30–35 rpm for approximately 3 min. After 2 h of incubation at room temperature, the unreacted dye is captured by the addition of 0.1 M L-cysteine and incubated for another 1 h at room temperature. The labeled vimentin is separated from free dye by size exclusion chromatography (Bio-Gel P-30, Bio-Rad Laboratories GmbH, Feldkirchen, Germany) using a column of size 1.0 × 30 cm (Econo-column, Bio-Rad). The column is filled with the bio-gel media and washed twice with the labeling buffer, before the addition of protein. After size exclusion chromatography, the protein concentration is quantified by UV absorption spectroscopy using an extinction coefficient of $\varepsilon_{\text{protein}}(280\,\text{nm}) = 24.87 \times 10^3$ $\text{M}^{-1}$ $\text{cm}^{-1}$ and a molecular mass of 53.88 kDa. The labeled protein is dialyzed overnight against storage buffer (2 mM PB with 8 M urea, pH 7.5) at 4°C. After dialysis, the labeling ratio is determined by measuring the protein concentration and amount of Atto647N bound to the protein using an extinction coefficient of $\varepsilon_{dye}(647\,\text{nm}) = 1.5 \times 10^5$ $\text{M}^{-1}$ $\text{cm}^{-1}$, and correction factor $cf(280\text{ nm}) = 0.05$. The degree of labeling is calculated as:

$$\text{labeling ratio} = \frac{A(647\,\text{nm})\varepsilon_{\text{protein}}(280\,\text{nm})}{(A(280\,\text{nm}) - A(647\,\text{nm})cf)\varepsilon_{\text{dye}}(647\,\text{nm})} \tag{1}$$

The labeled protein is aliquotted and stored at -80°C.

#### Functionalization of polystyrene beads with maleimide

Carboxylated polystyrene beads (PPs-4.2 COOH, 4.0–4.4 $\mu$m diameter, 5% w/v, Kisker Biotech, Steinfurt, Germany) are functionalized with $NH_2$-PEG-Maleimide and $NH_2$-PEG-OH following the protocol described in Refs.[6,7] The functionalized beads are stored in PBS with 2% BSA at 4°C until use.

#### Single filament interaction measurements

Interaction measurements are performed at room temperature using an optical trap (C-Trap, Lumicks, Amsterdam, The Netherlands) in quadruple trap mode as described in Refs.[8,9] The setup (Fig. 1a) combines a pressure driven microfluidic flow cell to flush the samples, and a confocal microscope with excitation lasers of wavelength 638 nm and 532 nm. The microfluidic flow cell has three inlets (Fig. 1b): subchannel (I) contains maleimide-functionalized polystyrene beads with a diameter of $\approx 4.2$ $\mu$m in assembly buffer, (II) contains assembly buffer (2 mM PB with 100 mM KCl, pH 7.5), and (III) contains 1 mg/L vimentin filaments in assembly buffer. During the measurements, the positions of the beads are recorded by particle tracking in bright-field images, while the filaments are observed with confocal microscopy. To start each interaction measurement, four beads are captured with the optical tweezers in subchannel (I). These beads are brought to the assembly buffer channel (II) to calibrate the trap stiffness. The trap stiffness is calibrated by the power spectral density of the thermal fluctuations of the trapped beads in subchannel (II) without flow. The beads are moved to subchannel (III) and a weak flow is applied. Single vimentin filaments are captured between each bead pair: filament F12 is tethered between beads b1 and b2 and filament F34 is tethered between beads b3 and b4. The traps with vimentin filaments are moved back to subchannel (II) and the flow is then stopped. We start with both filaments aligned in parallel within the same $z$-plane. Filament F34 is then rotated by 90° relative to filament F12. Next, the beads associated with filament F34 are lowered to a position of -4.7 $\mu$m along the $z$-axis. To obtain a cross configuration, filament F12 is then adjusted so that it lies within the same $x$-$y$ plane as the filament F34. Throughout this positioning procedure, bright-field images of the beads are used to guide the alignment. After F12 is positioned, filament F34 is raised back to the original $z$-plane so that it crosses filament F12. Since filaments behave as entropic springs, the filaments are pre-stretched to 2 pN to reduce thermal fluctuations, allowing them to straighten and interact in the same plane. The forces applied by all optical traps are then set to 0 pN. The filament F34 is moved back and forth in the $y$-direction at different pulling velocity conditions $v = 0.05$ $\mu$m/s, 0.1 $\mu$m/s, 0.5 $\mu$m/s, 1 $\mu$m/s, and 5 $\mu$m/s. The force acting on b1 is recorded for data analysis, as our experimental setup allows for direct measurement of force only from bead b1, and not from b2.

#### Data analysis

Force and time data are recorded at a sampling frequency of 78 kHz. To standardize the analysis across all pulling velocity conditions, the raw data are downsampled to 100 Hz using the `downsampled_by()` function from the Lumicks Pylake package.[10] Downsampling is performed by dividing the raw data into consecutive non-overlapping blocks of 780 points and replacing each block with their mean value. A sampling rate of 100 Hz remains sufficient to resolve the relevant force dynamics with reduced computational cost. After downsampling, the portion of the force–time data containing an interaction is manually selected. An interaction is defined as the interval in which the force deviates from the baseline, starting with an initial rise in force and ending with a rupture event marked by a rapid force drop (Fig. 2a(i)). A small linear force offset is observed in some datasets due to coupling between the energy potentials of the optical traps.[9] To remove this offset, a baseline correction is performed. First, the boundaries of the interaction are detected: the force signal is smoothed using two moving averages with different window sizes $w$ (50 and 400 data points, Fig. 2a(ii)). The difference between the two smoothed signals is normalized by the mean of the first 50 points of the 400-point smoothed signal and the absolute value of this normalized difference is reported as a percentage. The start of the interaction is identified as the first time point at which this difference exceeds a threshold of $0.8\sigma$, where $\sigma$ is the standard deviation of the first 50 force measurements (Fig. 2a(iii)). The end of the interaction is defined as the time point of the peak force, corresponding to the rupture event. The flat non-interacting regions before the start and at the end of the trajectory (red scatter, Fig. 2a(iv)) are then used to estimate a baseline via linear regression, which is subsequently subtracted from the force signal. A geometry correction factor, derived in Refs.,[8,9] is applied to obtain the force acting at the interaction site. Since only the force on bead b1 can be directly measured, and forces along the x-axis are balanced ($F_{1x} = F_{2x}$), $F_{2y}$ can be recovered from the filament geometry alone. The total interaction force $F = F_{1y} + F_{2y}$ is then given as:

$$F = F_{1y} cf_{\text{geo}} \tag{2}$$

where $F_{1y}$ is the measured force on bead b1. The correction factor $cf_{\text{geo}}$ is defined as:

$$cf_{\text{geo}} = \frac{l_{12} + a}{l_2 + a/2}, \tag{3}$$

where $a$ is the bead diameter; $l_1$ and $l_2$ are the distances from beads b1 and b2 to the vertical filament, with $l_{12} = l_1 + l_2$. Although the symmetric case ($l_1 = l_2$, Fig. 2b(ii)) is typical, $l_1$ and $l_2$ are measured independently to handle asymmetric configurations (Fig. 2b(iii)). The calculation of $cf_{\text{geo}}$ is automated by determining the intersection of the two filaments based on the bead positions. As the bead positions are recorded at 4 Hz using bright-field imaging, the bead position data is linearly interpolated using `numpy.linspace` to match the temporal resolution of the downsampled force data. Interactions with rupture forces below 1 pN are excluded, as such low forces could result from noise rather than binding events. The resulting force–time trajectories are used for Bayesian inference.

### Bayesian inference of bond parameters

#### Bayesian inference

We model the interaction between two vimentin filaments as a single bond using Bell-Evans kinetics.[11,12] This is a two-state model where the filaments can be either in a bound state or unbound state (see Fig. 1 in main text). The force-dependent unbinding rate is given as:

$$r(t) = r_u e\left(\frac{x_u F(t)}{k_B T}\right) \tag{4}$$

where $r_u$ is the force-independent unbinding rate, $x_u$ is the distance from the bound state to the transition state, and $F(t)$ is the force acting on the bond. The goal of our Bayesian approach is to infer the underlying bond parameters, $r_u$ and $x_u$, from experimental force-time trajectories $F(t)$. For each data point in the experimental force–time trajectory, we record the measured force $F_i$ and assign it to either a bound or unbound state. The corresponding state label is denoted $S_i$, with $S_i = 1$ for the bound state and $S_i = 0$ for the unbound state. We refer to the full set of observations $\{F_i, S_i\}$ simply as the data.

Our objective is to compute the posterior probability distribution $P(r_u, x_u|\{F_i, S_i\})$, which quantifies how plausible the bond parameter pairs ($r_u$, $x_u$) are given the observed data. By Bayes' theorem, the posterior distribution is given by:

$$P(r_u, x_u|\{F_i, S_i\}) = \frac{P(\{F_i, S_i\}|r_u, x_u)P(r_u, x_u)}{P(\{F_i, S_i\})} \tag{5}$$

where $P(r_u, x_u)$ is the prior, $P(\{F_i, S_i\}|r_u, x_u)$ is the likelihood, and $P(\{F_i, S_i\})$ is the evidence. The evidence is computed by marginalizing over all possible parameter values:

$$P(\{F_i, S_i\}) = \int_{r_u}\int_{x_u} P(\{F_i, S_i\}|r_u, x_u)P(r_u, x_u)\mathrm{d}r_u\mathrm{d}x_u \tag{6}$$

and therefore does not depend on any particular choice of $r_u$ and $x_u$. The evidence acts as a normalization constant that guarantees the posterior is a valid probability distribution, but depends on the choice of model and prior. Additionally, computing the evidence requires a high-dimensional integral over the entire parameter space, which is computationally intractable for most models. Thus for posterior evaluation we only focus on the prior distribution and the likelihood:

$$P(r_u, x_u | \{F_i, S_i\}) \propto P(\{F_i, S_i\} | r_u, x_u) P(r_u, x_u) \tag{7}$$

This proportionality is sufficient for parameter estimation as Markov Chain Monte Carlo (MCMC) sampling methods[13] only evaluate relative probabilities.

The likelihood is the conditional probability of observing the data given bond parameter pairs $(r_u, x_u)$. Since the Bell-Evans model[11,12] is a two-state system in which the bond exists in one of two states, and transitions between states are governed by force-dependent unbinding rate, $r(t)$, the likelihood for an individual data point $(F_i, S_i)$ follows a Bernoulli distribution which can be written as:

$$P(F_i, S_i | r_u, x_u) = \begin{cases} e^{-r(t_i)\mathrm{d}t_i} & \text{if } S_i = 1 \\ 1 - e^{-r(t_i)\mathrm{d}t_i} & \text{if } S_i = 0 \end{cases} \tag{8}$$

where $S_i = 1$ corresponds to bond survival and $S_i = 0$ to an unbinding event. Equation (8) arises because over the time interval $dt_i$, the probability of bond survival decays exponentially with a detachment rate $r(t_i)$.

For $N$ observations, the full likelihood can be written as the product of the likelihood of the individual data points. This is because of our assumption that it is a Markovian process (the bond has no memory), where the probability of a transition in a time interval $dt_i$ depends only on the current bond state $S_i$ and the instantaneous force $F_i$, and is independent of the history.

For this inference, we use a flat prior where all values of $r_u$, $x_u$ are equally likely in some parameter range. We use broad uniform priors over the ranges $r_u \in \{0, 1\}$ (in $\mathrm{s}^{-1}$) and $x_u \in \{0, 1\}$ (in nm) which are much wider than the resulting posteriors.

#### Censoring correction

In the optical tweezer measurements, we only analyze trajectories with breaking forces that exceed 1 pN. Trajectories with breaking events below 1 pN cannot be reliably distinguished from noise and are therefore omitted. This introduces a censoring effect, the analyzed dataset $\mathcal{D}_{\text{obs}} = \{F_i, S_i\}_{i=1}^{N}$ contains only trajectories where the bond ruptured above 1 pN, whereas the true population also includes earlier unbinding events that remain unobserved. To correct for this selection bias, we incorporate a censoring term into the likelihood that accounts for the conditional survival of the bond up to the 1 pN threshold. For each observed trajectory $j$, we compute the probability that the bond survives all time points where the force is below the detection threshold $F_{threshold} = 1$ pN:

$$P(\text{survive}_j \mid r_u, x_u) = \prod_{F<F_{\text{threshold}}} e^{-r(t)\text{dt}} \tag{9}$$

Because $\mathcal{D}_{\text{obs}}$ contains only trajectories where the bond did survive until $F_{\text{threshold}}$, the likelihood must be renormalized by conditioning on this event. Therefore the posterior is written as:

$$P(r_u, x_u|\mathcal{D}_{\text{obs}}) \propto \frac{P(\mathcal{D}_{\text{obs}}|r_u, x_u)P(r_u, x_u)}{\prod_{j=1}^{M} P(\text{survive}_j|r_u, x_u)} \tag{10}$$

#### Posterior sampling

To characterize the posterior distribution, we use a Markov Chain Monte Carlo (MCMC) sampling[13], a widely used computational approach for exploring high-dimensional and complex probability distributions in Bayesian analysis. MCMC generates samples from the posterior by constructing a Markov chain whose stationary distribution matches the target posterior distribution. Given sufficient iterations, the MCMC samples provide an accurate discrete approximation of the posterior distribution. Specifically, we use the No-U-Turn Sampler (NUTS)[14], an advanced variant of Hamiltonian Monte Carlo[15] that exploits gradient information to efficiently explore the posterior landscape. We perform 4 independent chains, each with 2,000 warm-up iterations and 5,000 posterior draws, yielding 20,000 total posterior samples. The effective sample size for all parameters exceed 4600. Convergence is assessed using the $\hat{R}$ diagnostic[16] and by visual inspection of trace plots. All sampling is performed using the PyMC probabilistic programming library[17,18].

#### Posterior visualization

To visualize the posterior of bond parameter pairs $(r_u, x_u)$, the posterior samples from PyMC are converted into a continuous probability density using two-dimensional Gaussian kernel density estimation (KDE), implemented via `scipy.stats.gaussian_kde`. The KDE provides a smooth approximation of the posterior density by placing a Gaussian kernel at each sample point and summing the contributions. The resulting density is evaluated on a 200 $\times$ 200 grid spanning the parameter space and normalized such that the maximum density equals 1. We plot only the 95% credible region, defined as the smallest region of parameter space containing 95% of the posterior probability.

#### Information gain

To quantify the information gain from prior to posterior, we use the Kullback-Leibler (KL) divergence[19]. Let $q \equiv q(r_u, x_u \mid \mathcal{D}_{\text{obs}})$ denote the joint posterior and $p \equiv p(r_u)p(x_u)$ the factorized uniform prior. The KL divergence is:

$$D_{KL}(q \| p) = \mathbb{E}_q\left[\log \frac{q}{p}\right] \approx \frac{1}{N}\sum_{i=1}^{N} \log \frac{q_i}{p_i} \tag{11}$$

The joint KL divergence decomposes as:

$$D_{KL}(q \| p) = D_{KL}(q_r \| p_r) + D_{KL}(q_x \| p_x) + I(r_u; x_u) \tag{12}$$

where $q_r \equiv q(r_u \mid \mathcal{D}_{\text{obs}})$, $q_x \equiv q(x_u \mid \mathcal{D}_{\text{obs}})$, and $I(r_u; x_u)$ is the mutual information capturing posterior correlations between parameters. The first two terms quantify the information gained about $r_u$ and $x_u$ individually.

Since the prior is uniform on $[0, 1]$, $\log p_r = \log p_x = 0$, the KL divergences reduce to expectations of the log posterior density. The joint posterior and marginals are approximated by KDEs fitted to the MCMC samples, giving the Monte Carlo estimates:

$$D_{KL}(q_r \| p_r) \approx \frac{1}{N} \sum_{i=1}^{N} \log \hat{q}_r(r_u^{(i)}) \tag{13}$$

$$D_{KL}(q_x \| p_x) \approx \frac{1}{N} \sum_{i=1}^{N} \log \hat{q}_x(x_u^{(i)}) \tag{14}$$

$$I(r_u; x_u) \approx \frac{1}{N} \sum_{i=1}^{N} \log \frac{\hat{q}(r_u^{(i)}, x_u^{(i)})}{\hat{q}_r(r_u^{(i)})\, \hat{q}_x(x_u^{(i)})} \tag{15}$$

All values are reported in bits.

### References


(1) Herrmann, H.; Kreplak, L.; Aebi, U. *Methods in Cell Biology*; Elsevier, 2004; Vol. 78; pp 3–24.

(2) Forsting, J.; Kraxner, J.; Witt, H.; Janshoff, A.; Köster, S. Vimentin Intermediate Filaments Undergo Irreversible Conformational Changes during Cyclic Loading. *Nano Letters* **2019**, *19*, 7349–7356.

(3) Winheim, S.; Hieb, A. R.; Silbermann, M.; Surmann, E. M.; Wedig, T.; Herrmann, H.; Langowski, J.; Mücke, N. Deconstructing the late phase of vimentin assembly by total internal reflection fluorescence microscopy (TIRFM). *PLoS ONE* **2011**, *6*, e19202–e19202.

(4) Nöding, B.; Köster, S. Intermediate filaments in small configuration spaces. *Physical Review Letters* **2012**, *108*, 088101.

(5) Block, J.; Witt, H.; Candelli, A.; Peterman, E. J.; Wuite, G. J.; Janshoff, A.; Köster, S. Nonlinear Loading-Rate-Dependent Force Response of Individual Vimentin Intermediate Filaments to Applied Strain. *Physical Review Letters* **2017**, *118*, 048101.

(6) Janissen, R.; Berghuis, B. A.; Dulin, D.; Wink, M.; Van Laar, T.; Dekker, N. H. Invincible DNA tethers: Covalent DNA anchoring for enhanced temporal and force stability in magnetic tweezers experiments. *Nucleic Acids Research* **2014**, *42*, e137.

(7) Block, J.; Witt, H.; Candelli, A.; Danes, J. C.; Peterman, E. J. G.; Wuite, G. J. L.; Janshoff, A.; Köster, S. Viscoelastic properties of vimentin originate from nonequilibrium conformational changes. *Science Advances* **2018**, *4*, eaat1161.

(8) Schepers, A. V.; Lorenz, C.; Nietmann, P.; Janshoff, A.; Klumpp, S.; Köster, S. Multiscale mechanics and temporal evolution of vimentin intermediate filament networks. *Proceedings of the National Academy of Sciences* **2021**, *118*, e2102026118.

(9) Schaedel, L.; Lorenz, C.; Schepers, A. V.; Klumpp, S.; Köster, S. Vimentin intermediate filaments stabilize dynamic microtubules by direct interactions. *Nature Communications* **2021**, *12*, 3799.

(10) Vanlier, J.; Pauszek, R.; Moldovan, D.; van den Berg, A.; Jachowski, T.; Mirone, A.; Broekmans, O.; Moerland, R.; Moyo, A.; lilfer; Lamerton, S. lumicks/pylake: v1.8.0. 2025; `https://zenodo.org/doi/10.5281/zenodo.4280788`.

(11) Bell, G. I. Models for the Specific Adhesion of Cells to Cells: A theoretical framework for adhesion mediated by reversible bonds between cell surface molecules. *Science* **1978**, *200*, 618–627.

(12) Evans, E.; Ritchie, K. Dynamic strength of molecular adhesion bonds. *Biophysical Journal* **1997**, *72*, 1541–1555.

(13) Gelman, A.; Carlin, J. B.; Stern, H. S.; Dunson, D. B.; Vehtari, A.; Rubin, D. B. *Bayesian Data Analysis*; Chapman and Hall/CRC, 2013.

(14) Hoffman, M. D.; Gelman, A. The No-U-Turn Sampler: Adaptively Setting Path Lengths in Hamiltonian Monte Carlo. *Journal of Machine Learning Research* **2014**, *15*, 1593–1623.

(15) Duane, S.; Kennedy, A.; Pendleton, B. J.; Roweth, D. Hybrid Monte Carlo. *Physics Letters B* **1987**, *195*, 216–222.

(16) Gelman, A.; Rubin, D. B. Inference from Iterative Simulation Using Multiple Sequences. *Statistical Science* **1992**, *7*, 457–472.

(17) Salvatier, J.; Wiecki, T. V.; Fonnesbeck, C. Probabilistic programming in Python using PyMC3. *PeerJ Computer Science* **2016**, *2*, e55.

(18) Abril-Pla, O.; Andreani, V.; Carroll, C.; Dong, L.; Fonnesbeck, C. J.; Kochurov, M.; Kumar, R.; Lao, J.; Luhmann, C. C.; Martin, O. A.; Osthege, M.; Vieira, R.; Wiecki, T.; Zinkov, R. PyMC: a modern, and comprehensive probabilistic programming framework in Python. *PeerJ. Computer Science* **2023**, *9*, e1516.

(19) Cover, T. M.; Thomas, J. A. *Elements of information theory*; Wiley series in telecommunications; Wiley: New York, 1991; Chapter 2.

#### Supplementary figures

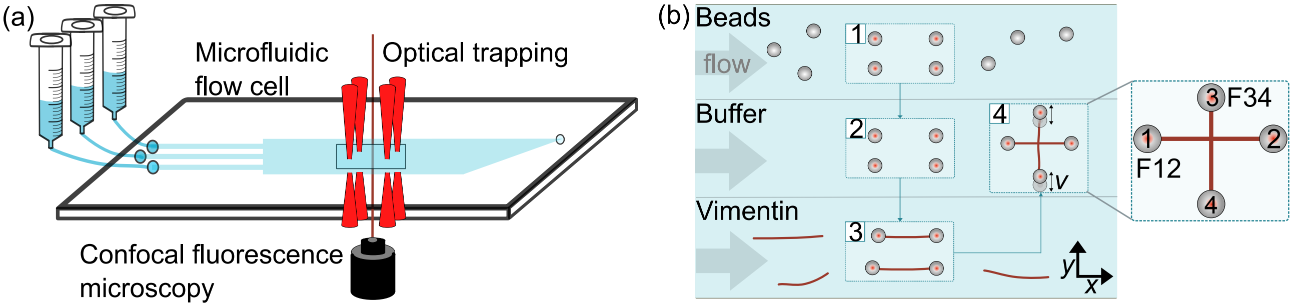


Figure 1: (a) Schematic of the quadruple optical trap setup with confocal microscopy and a microfluidic device for measuring interactions between single filaments. (b) Schematic of the microfluidic flow cell: (1) Four beads are captured using optical traps. (2) These beads are transferred to the measurement buffer for trap calibration. (3) The beads are moved to the vimentin channel and a single vimentin IF is tethered between each pair of beads. F12 is the filament between beads 1 and 2 and F34 is the filament between beads 3 and 4. (4) The two tethered vimentin IFs are brought into contact in a crossed configuration, and the vertical vimentin IF is moved back and forth in the $y$-direction.

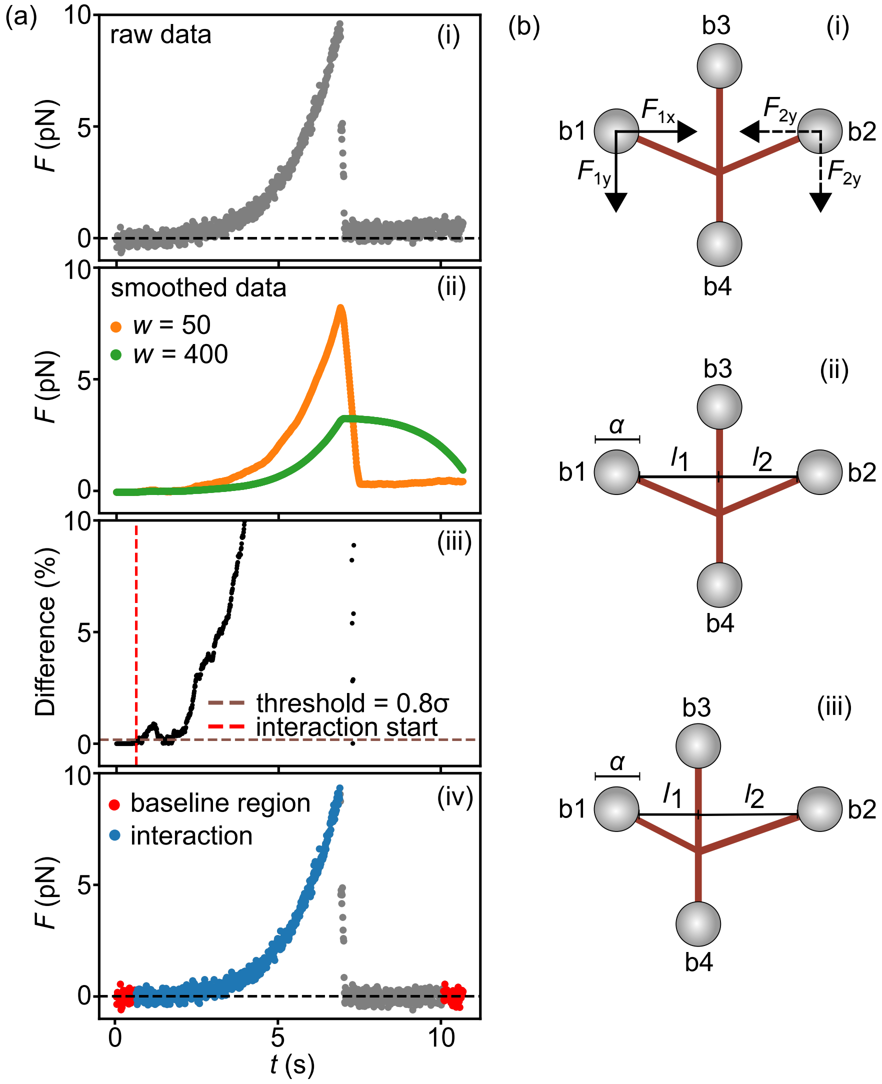


Figure 2: (a) Illustration of the data processing pipeline for a representative force-time trajectory. (i) The interaction region is manually selected as the interval between the initial force rise and the rupture event. (ii) The force signal is smoothed with two moving averages of different window sizes $w$ (50 and 400 data points). The difference between the two smoothed signals is normalized by the mean of the first 50 points of the larger-window signal, and the absolute value of this quantity is used to detect interaction boundaries. (iii) The start of the interaction is identified as the first time point where the difference in percentage exceeds $0.8\sigma$, where $\sigma$ is the standard deviation of the first 50 force measurements; the end is defined as the time point of peak force before rupture. (iv) Force-time trajectory after baseline correction, with the baseline region (red scatter) and interaction event (blue scatter) indicated. (b) (i) Schematic of the crossed-filaments experiment. The force components $F_{1x}$ and $F_{1y}$ on bead b1 (solid arrows) are directly measured, while $F_{2x}$ and $F_{2y}$ on bead b2 (dashed arrows) are not accessible experimentally. (ii) Symmetric case ($l_1 = l_2$), where $l_1$ and $l_2$ are the distances from beads b1 and b2 to the vertical filament and $a$ is the bead diameter. These geometric quantities enter $cf_{\text{geo}}$ and are used together with $F_{1y}$ to obtain the total force at the interaction site. (iii) Asymmetric case ($l_1 < l_2$), illustrating that the same geometric quantities and force balance apply when the distances differ, as accounted for by measuring $l_1$ and $l_2$ independently.[8,9]

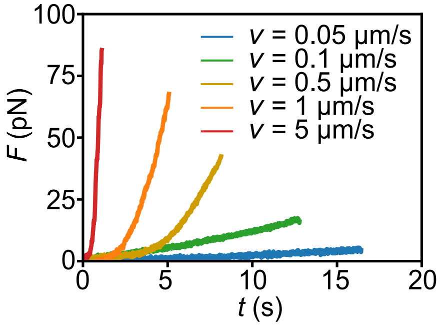


Figure 3: Representative force-time curves at each pulling velocity. At $v = 5$ $\mu$m/s force builds rapidly, causing the bond to rupture quickly. At $v = 0.05$ $\mu$m/s, force increases slowly, exposing the bond to lower stress over a longer time before rupture. The resulting distributions of breaking forces are shown in Fig. 3a in the main text.

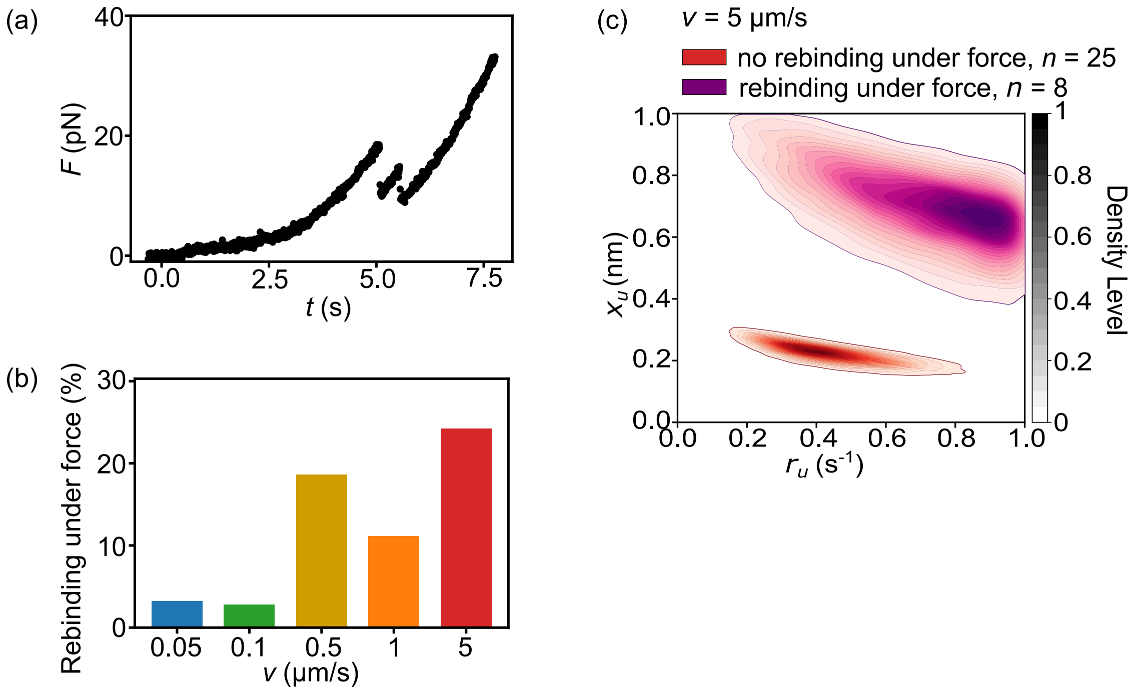


Figure 4: (a) Representative force-time curve showing rebinding events under force. (b) The frequency of events with rebinding under force increases with pulling velocity $v$. (c) The posterior distributions of rebinding events under force (purple) are inconsistent with those obtained from events without rebinding under force (red). These rebinding events are excluded from the analysis.

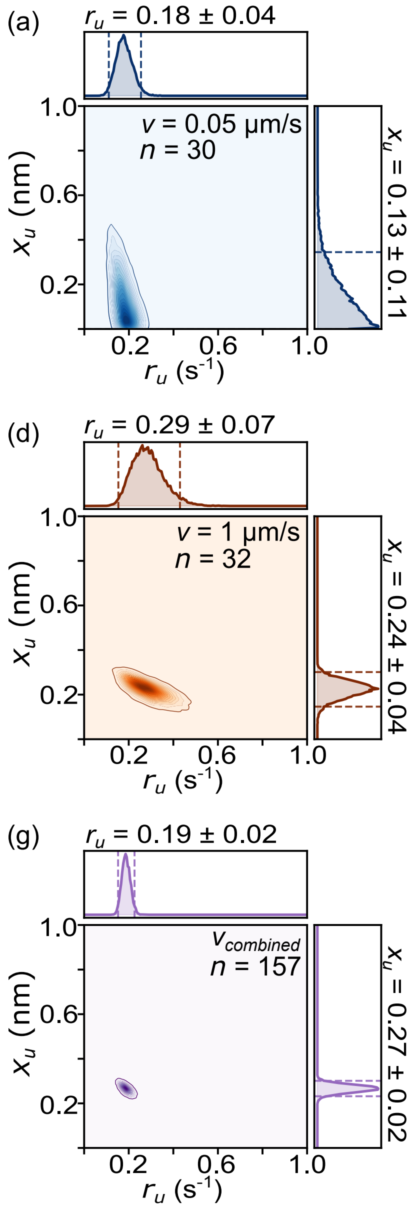


Figure 5: (a) Joint posterior distributions of $r_u$ and $x_u$ for each pulling velocity $v$. The 2D contour plots show the joint posterior, with marginal distributions of $r_u$ (top) and $x_u$ (right), the light shaded region spanning the full parameter range indicates the uniform prior. Dashed lines indicate the 95% highest density interval. The mean and standard deviations of each parameter are annotated on the respective marginal distributions.

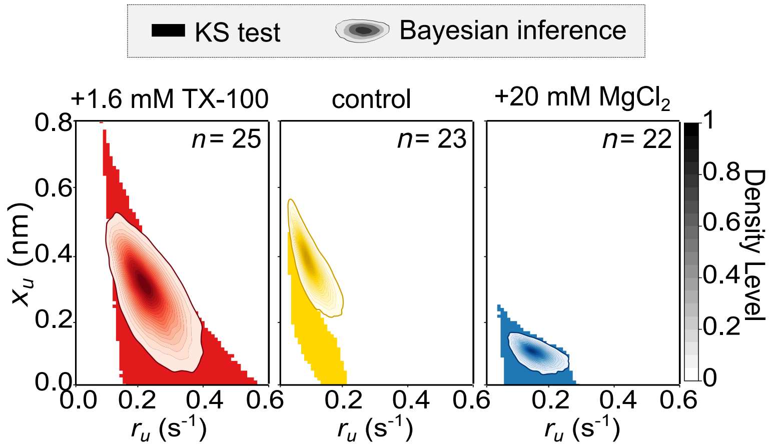


Figure 6: Application of the Bayesian inference framework to previously obtained MT–vimentin interaction data using different buffers[9] demonstrates the generalizability of the method for different cytoskeletal filament interactions.
